# Scale-dependent spatial organisation of action observation network responses to hand actions and emotional faces

**DOI:** 10.64898/2026.07.28.741155

**Authors:** Yusuke Haruki, Risako Yamaguchi, Kenji Ogawa

## Abstract

Human social perception relies on multiple visual signals, including object-directed actions and emotional facial expressions. It remains unclear whether responses to these signals converge within action-observation regions and whether apparent convergence depends on the spatial scale of analysis. We scanned 39 healthy human adults during two block-design fMRI tasks, hand action observations contrasted with paper-tube controls, and emotional expression observations contrasted with neutral faces. Using independently defined meta-analytic action observation network (AON) ROIs in the inferior frontal gyrus (IFG), inferior parietal lobule (IPL), and posterior superior temporal sulcus (pSTS), we analysed responses at whole-brain, ROI-average, and AON-constrained voxelwise scales. Whole-brain overlap between the two contrasts was localised primarily to the left lateral occipitotemporal extrastriate cortex, outside the AON. At the regional ROI-average scale, the IFG showed positive responses to both contrasts, whereas the IPL and pSTS showed opposing task preferences. Crucially, regional IFG co-engagement did not imply that the same voxels responded to both contrasts. Suprathreshold voxelwise overlap within the AON was virtually absent, and individual-level analysis revealed a small but reliable posterior displacement of hand action relative to emotional face within IFG. These findings show that spatial convergence between hand and emotion observation responses depends on the scale of analysis. The same IFG region can be engaged by both stimulus domains while retaining distinct local response topographies.

## Introduction

Seeing another person act on the world and seeing another person express emotion are both core problems for social perception. The action observation network (AON) is commonly described as a distributed set of frontal, parietal, and temporal regions involved in observing and interpreting others’ actions. Classical mirror neuron accounts and meta-analytic work have repeatedly implicated inferior frontal gyrus (IFG), inferior parietal lobule (IPL), and posterior superior temporal sulcus (pSTS) in action observation, imitation, and related sensorimotor tasks (Caspers et al., 2010; Hardwick et al., 2018; Molenberghs et al., 2012; Rizzolatti & Craighero, 2004). Work on observed object-directed hand actions further indicates that parietal and premotor regions carry action information beyond low-level visual variation (Ogawa & Inui, 2011). However, the individual subregions of the AON may not function as a single unit during social perception. Other social cues, such as emotional faces, might activate the frontal, parietal, and temporal areas differently than object-directed actions.

Facial expressions are closely linked to sensorimotor and affective accounts of emotion perception. Observing or imitating emotional facial expressions can engage the IFG and limbic systems (Carr et al., 2003; Leslie et al., 2004), and broader reviews emphasise sensorimotor contributions to facial expression perception (Wood et al., 2016). At the same time, emotional faces recruit visual and affective systems in addition to regions associated with action observation (Barrett et al., 2019; Haxby et al., 2000; Pitcher & Ungerleider, 2021). Recent causal work further supports pSTS-centred circuits in emotional expression perception, including recurrent interactions between pSTS and early visual cortex (Borgomaneri et al., 2023). Therefore, activation within the AON during face observation does not necessarily imply that action and face observation rely on the same underlying neural substrate. Rather, it may reflect partial spatial convergence between neural systems supporting object-directed action and facial expression observations.

A key challenge is that apparent spatial convergence can vary across scales of analysis. That is, whether convergence is assessed as overlap between whole-brain activations or regional signals within regions of interest (ROIs) might draw different conclusions (Morrison & Downing, 2007). This distinction is relevant to neural reuse, in which the same cortical region plays roles in multiple functions (Anderson, 2010). Many claims of a “shared” social network rely on group-level activation maps or ROI averages (Keysers & Gazzola, 2009; Molenberghs et al., 2012; Schmidt et al., 2021). Effective connectivity work also suggests that social cognitive recruitment of the human mirror system is organised through interactions among the STS, IPL, and IFG, rather than through a single frontoparietal module (Sadeghi et al., 2022). However, macro-level regional coactivation does not guarantee shared neural representations at a finer spatial scale. A single anatomical region can support distinct functions through spatially biased or distributed response patterns (Dinstein et al., 2008). Evaluating the spatial relationship between action– and face-related activations across different scales can help clarify whether regional engagement by both tasks reflects voxel-level overlap or a more local topographic organisation.

The present study directly investigated this scale-dependent mapping question using fMRI, where participants completed a hand action observation task and an emotional face observation task. Using bilateral IFG, IPL, and pSTS regions defined independently from meta-analytic coordinates (Caspers et al., 2010), we compared the two tasks across three spatial scales. Specifically, we mapped whole-brain overlap relative to AON boundaries, evaluated ROI-average profiles to test for regional engagement by both tasks, and analysed the voxel-level distribution of these activations within the AON. This approach allowed us to ask whether apparent convergence reflects the same-voxel overlap or a local topographic bias within the same region.

## Materials and methods

### Participants

Thirty-nine healthy adults (27 men, 12 women; mean age = 21.90 years, SD = 2.14; age range = 19–28 years) took part in the present study. All participants were native Japanese and right-handed as assessed by the Japanese version of the Edinburgh Handedness Inventory (Hatta & Nakatsuka, 1975; Oldfield, 1971). They reported normal or corrected-to-normal vision. Exclusion criteria were a history of neurological disorder, current psychotropic medication use, and standard MRI contraindications. Participants were recruited from Hokkaido University through campus advertisements, online bulletin boards, and word of mouth. Upon arrival at the experiment room, participants received a verbal and written explanation of the study procedures, potential risks, and their rights as research participants. All participants provided written informed consent and received 4000 JPY for a single 2-hour session. The study protocol was approved in advance by the Ethics Committee of the Center for Experimental Research in Social Sciences at Hokkaido University, and all procedures were conducted in accordance with the latest version of the Declaration of Helsinki.

### Task design

Participants completed two observation tasks during a single scanning session. Each task used a block-design with 12 s active blocks alternating with 12 s fixation periods, which served as implicit baseline. The motor observation task was always administered first, followed by the face observation task. This order was fixed a priori to minimise possible carryover from emotional face exposure to the motor observation run (Del Vecchio et al., 2024).

The motor observation task was designed to examine brain activity during the visual observation of object-directed hand actions involving everyday objects, contrasted with a visually matched non-biological control condition (Moriguchi et al., 2009) (Figure 1A). Stimuli were short colour video clips depicting common objects, such as cups, forks, and oranges, being approached and touched by either a human hand, in the action condition, or a paper tube, in the control condition. For the control condition, the paper tube was matched in size, shape, and approach trajectory to the human hand. Videos were presented at 1280 × 720 pixels and centred on a grey background. Each 12 s block consisted of four 3 s clips. Video clips were recorded by the experimenter using a digital camera under constant lighting and viewing angle conditions. The run contained 12 hand action and 12 paper control blocks, presented in alternating order with the starting condition fixed as paper control first. To ensure continuous attention, one clip in half of the blocks featured a change in approach direction, which participants detected via button press with their right index finger.

**Figure 1.**
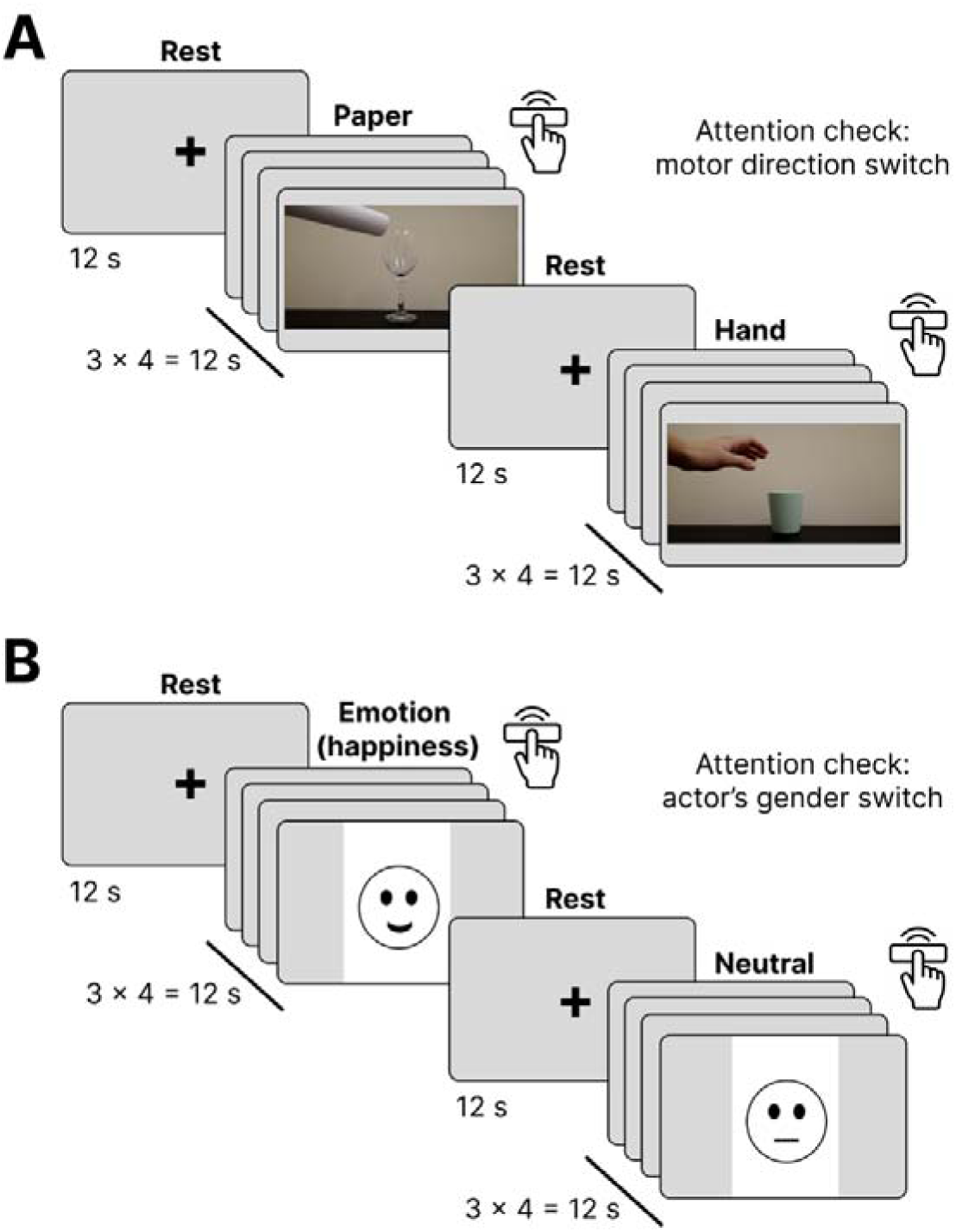
Task design schematic. Participants completed two block-design observation tasks. (A) In the motor observation task, participants viewed short video clips of everyday objects being approached and touched by either a human hand or a paper tube. Each active block lasted 12 s and contained four 3 s clips. Paper control and hand action blocks alternated with 12 s fixation periods. Participants detected occasional changes in motor approach direction. (B) In the face observation task, participants viewed static emotional or neutral faces from the AIST Face Database. Each active block lasted 12 s and contained four 3 s images. Emotional and neutral face blocks alternated with 12 s fixation periods. Participants detected occasional changes in the actor’s gender. In accordance with the use conditions of the database, the facial images shown in this schematic are placeholders and do not reproduce the actual database stimuli.

In the face observation task, participants viewed static facial expressions from the AIST Face Database (Fujimura & Umemura, 2018). The stimulus set included four female and four male actors displaying neutral expressions or one of six emotional expressions, anger, disgust, fear, happiness, sadness, and surprise (Figure 1B). For anger and disgust, open-mouth expressions were used. All non-neutral expressions were collapsed into a single emotional face condition. Each active block contained four 3 s images of the same emotional category, displayed at 750 × 1000 pixels. A total of 12 emotional face and 12 neutral face blocks were presented in alternating order with the starting condition of emotional face observation. As an attention check, one image in half of the blocks of each condition differed in gender from the others, which participants detected by pressing a button with their right index finger.

### MRI Acquisition

All MRI data were collected at Hokkaido University using a 3-Tesla Siemens Prisma scanner (Siemens, Erlangen, Germany) equipped with a 64-channel head coil. Participants lay supine in the scanner with their head immobilised using foam padding and a stabilising strap to minimise motion artefacts. Earplugs were provided to attenuate scanner noise. Visual stimuli were presented on a translucent rear-projection screen located at the back of the bore and viewed via an angled mirror mounted on the head coil. The experimental scripts were synchronised with the MRI data acquisition via TTL pulses, ensuring precise temporal alignment between stimulus presentation and image acquisition. Functional imaging was performed using a T2*-weighted gradient-echo echo-planar imaging (EPI) sequence sensitive to blood oxygenation level-dependent (BOLD) contrast. For each functional run, a total of 291 volumes were obtained and the first three volumes were discarded to allow for T1 equilibrium. Acquisition parameters were: repetition time (TR) = 2000 ms, echo time (TE) = 30 ms, flip angle = 90°, acquisition matrix = 94 × 94, in-plane resolution = 2.04 × 2.04 mm, 32 axial slices, slice thickness = 3.5 mm, inter-slice gap = 0.875 mm. These parameters provided whole-brain coverage. Anatomical images were acquired using a high-resolution T1-weighted magnetisation-prepared rapid acquisition gradient echo (MP-RAGE) sequence for spatial normalisation and coregistration. Parameters were: TR = 2300 ms, TE = 2.41 ms, inversion time (TI) = 900 ms, flip angle = 8°, 224 sagittal slices.

### Preprocessing

All preprocessing was performed with fMRIPrep version 24.1.1 (Esteban et al., 2019). The T1-weighted image underwent intensity non-uniformity correction, skull stripping, and tissue segmentation; spatial normalisation to MNI152NLin2009cAsym space used nonlinear registration with antsRegistration. For each BOLD run, head-motion correction and coregistration to the T1-weighted anatomy were performed with boundary-based registration. We used the preprocessed BOLD time series in MNI space and the corresponding brain mask for modelling. Before individual-level modelling, each run was converted to percent signal change within the brain mask. For each voxel, the post-dummy time series was divided by its run mean, multiplied by 100, and centred so that individual-level beta estimates were expressed in percent-signal-change (PSC) units. This scaling was used because the primary analyses compared contrast estimates across the two task runs.

### Individual-level GLM analysis

Statistical analyses were implemented in MATLAB using SPM25 functions (Tierney et al., 2025) for spatial smoothing, statistical distribution functions, and group-level modelling. Each run was modelled separately for each participant. For the main analysis, the preprocessed BOLD time series was spatially smoothed with a 6 mm FWHM Gaussian kernel before individual-level modelling. Task blocks were modelled as 12 s boxcars convolved with a canonical hemodynamic response function. The motor observation run included hand action and paper control regressors. The face observation run included emotional face and neutral face regressors. Nuisance regressors included the six rigid-body motion parameters, their first derivatives, framewise displacement, standardised DVARS, CSF signal, white-matter signal, all fMRIPrep cosine regressors, and binary fMRIPrep motion-outlier regressors. The binary spike regressors were the fMRIPrep motion_outlier columns; continuous nuisance regressors, including framewise displacement and standardised DVARS, were median-filled when necessary and standardised, whereas spike regressors were retained as binary columns. The fMRIPrep cosine regressors served as the high-pass component, and no additional temporal high-pass filter was applied. Button presses were modelled as zero-duration nuisance events when present. Blocks with button press omissions were excluded from the relevant condition regressors and included as invalid-block nuisance regressors. Across all participants and both tasks, 15 of 1872 planned condition blocks were marked invalid, leaving 1857 valid condition blocks for the primary condition regressors. Voxelwise individual-level models were fit to percent-signal-change time series. We created the primary individual-level contrasts as hand action vs. paper control and emotional face vs. neutral face.

### Whole-brain group analysis

Group level maps were generated using one sample t-tests for the positive hand action contrast and the positive emotional face contrast. For visualisation and overlap counting, maps were thresholded at voxelwise *p* < .001 uncorrected, with clusters of at least 15 voxels. The thresholded maps were then intersected to quantify whole-brain spatial overlap between the two contrasts.

### AON ROI-averaged activation analysis

We operationally defined the AON as an independent meta-analytic action-observation search space derived from Caspers et al. (2010). Six 8 mm radius spherical ROIs were created in MNI152NLin2009cAsym space: bilateral IFG (left: x = –50, y = 9, z = 30, 120 voxels; right: x = 52, y = 12, z = 26, 123 voxels), bilateral IPL (left: x = –60, y = –24, z = 36, 115 voxels; right: x = 44, y = –34, z = 44, 114 voxels), and bilateral pSTS (left: x = –54, y = –50, z = 8, 119 voxels; right: x = 56, y = –40, z = 4, 118 voxels). The full AON union contained 709 voxels. This search space was defined entirely independently of the present activation maps.

First, we extracted and averaged PSC estimates from the hand action > paper control and emotional face > neutral face contrasts from each of the six AON ROIs, for each participant. We then performed a three-way repeated measures ANOVA with the task (hand vs. paper action, emotional vs. neutral face), region (IFG, IPL, pSTS), and hemisphere (left, right) as within factors. Our primary interest was the task × region interaction, which evaluated whether the two tasks elicited distinct regional profiles. P-values for multiple comparisons were corrected using the false discovery rate (FDR) (Benjamini & Hochberg, 1995). Planned follow-up tests compared the task difference (i.e., hand vs. paper minus emotion vs. neutral) within each ROI family and within each unilateral ROI. The three ROI-family comparisons were treated as one FDR family, and the six unilateral ROI comparisons as a separate FDR family.

### Voxelwise topography in the AON ROI

To characterise the spatial relationship of the activations at the voxel level, we restricted the group-level t-maps to the independent AON search space. Overlap was assessed as the intersection of these maps at voxelwise *p* < .001 (uncorrected). For each region, we calculated voxel counts (for hand action only, emotional face only, and their overlap) and Dice coefficients.

Furthermore, we tested potential segregations for hand action and emotional face activations within the IFG. To this end, we computed t-weighted centre of gravity (COG) coordinates for each condition at the participant level. Unsmoothed individual-level t maps for each contrast (hand vs. paper, emotion vs. neutral) were restricted to the bilateral IFG ROIs separately and thresholded at height *p* < .05 (uncorrected, one-tailed). A participant contributed to a hemisphere if both contrasts elicited at least one positive suprathreshold voxel within that IFG ROI.

For each participant and hemisphere, the primary location summary was a t-weighted COG calculated as:

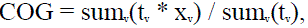

where t_v_ is the positive t-value at voxel v and x_v_ is its MNI coordinate vector. This statistic-weighted COG aligns with intensity-weighted COG implemented in FSL software (Jenkinson et al., 2012). Computing a weighted centroid within anatomical boundaries provides a robust localisation target that is less sensitive to noise and vascular displacement than peak coordinates.

To quantify the spatial offset between conditions, we computed the 3D displacement vector from the emotional face COG to the hand action COG per participant and hemisphere. These vectors were decomposed into lateral, anterior, and dorsal axes. We sign-flipped the left hemisphere’s x-axis so that positive values consistently indicate a more lateral, anterior, or dorsal hand action COG. Bootstrap 95% confidence intervals (CIs, based on 20,000 resamples with replacement across participants) summarised uncertainty in the displacement components and mean vector length. A one-sample Hotelling’s T² test evaluated whether the mean signed 3D displacement vector differed significantly from zero.

## Results

All 39 participants contributed to the analyses. Mean FD ranged from 0.050 to 0.285 mm, FD > 0.5 mm frame counts ranged from 0 to 38 per run, and the fMRIPrep motion-outlier regressors used in the GLM ranged from 0 to 206 per run. They performed the detection tasks for attention maintenance accurately. Mean block accuracy was.991 for hand action blocks,.987 for paper-control blocks,.981 for emotional face blocks, and .981 for neutral-face blocks. However, reaction times showed that the conditions were not perfectly matched for detection demand. Responses were slower for hand action than paper-control targets, mean difference = 0.191 s, 95% CI [0.164, 0.219], *t*(38) = 14.04, *p* < .001, Cohen’s d = 2.25. Responses were also slower for emotional face than neutral-face targets, mean difference = 0.095 s, [0.028, 0.161], *t*(38) = 2.90, *p* = .0062, d = 0.46. These behavioural data indicate high task engagement while also showing that the contrasts differed in detection demand.

### Whole-brain maps show overlap outside the AON

At group level, whole-brain thresholded maps showed distinct activation patterns for the two contrasts, along with a localised region of spatial overlap (Figure 2, Table 1). At height *p* < .001 (uncorrected) with clusters of at least 15 voxels retained, the hand action contrast (hand action > control) activated 2494 voxels. Dominant activation clusters were located in bilateral sensorimotor areas (postcentral/precentral gyrus), bilateral SPL, left premotor cortex, and left ventral occipitotemporal cortex (Table 1). Conversely, the emotional face contrast (emotional face > neutral face) activated 7161 voxels, with major clusters located bilaterally in occipitotemporal visual regions (including the fusiform gyrus), IFG, middle frontal gyrus, temporal poles, and the parahippocampal gyrus/amygdala region, as well as the medial prefrontal cortex (Table 1).

**Figure 2.**
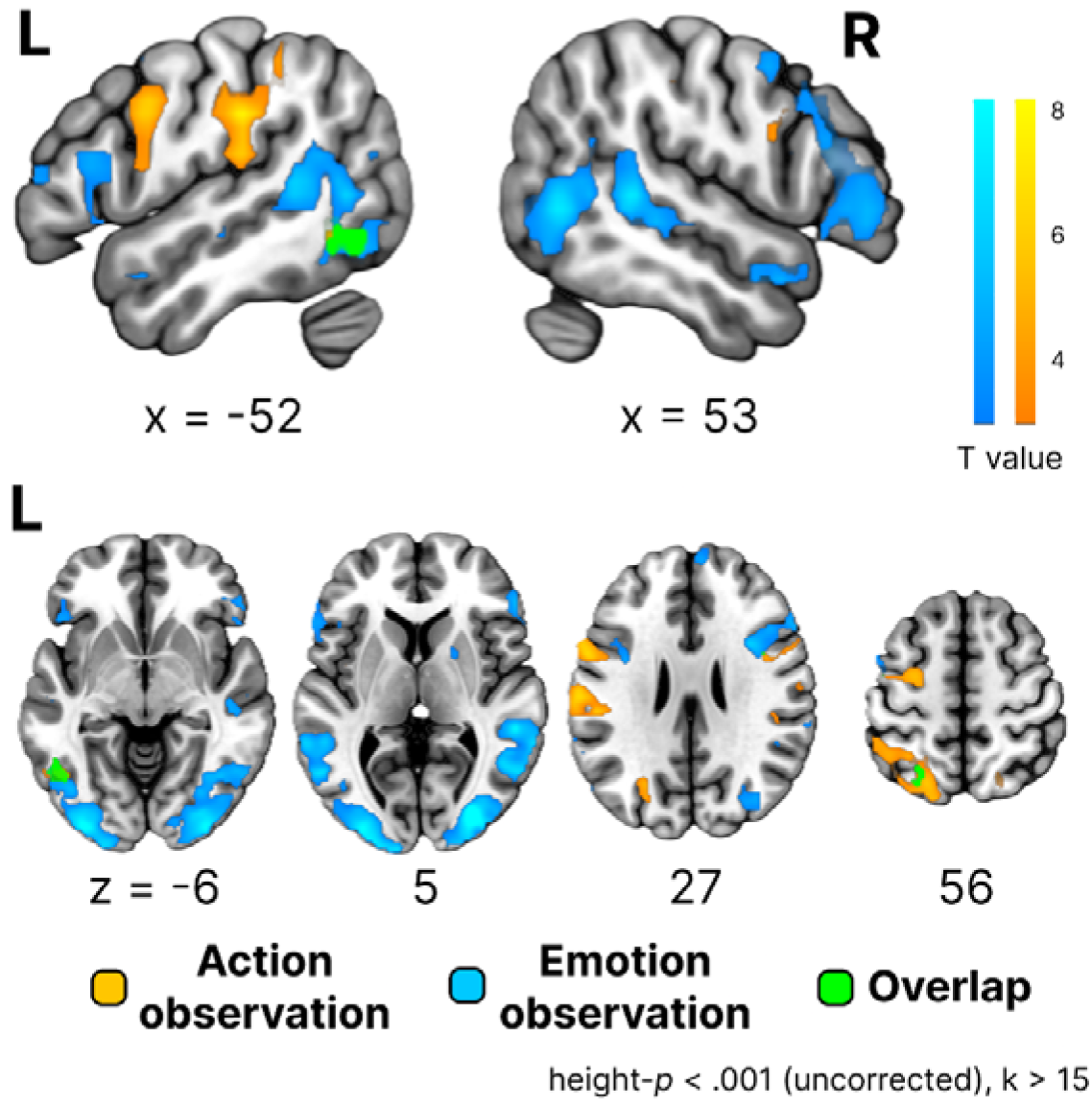
Whole-brain activation for hand action and emotional face observation. Group-level maps are shown for the hand action > paper control contrast and the emotional face > neutral face contrast. Maps were thresholded at voxelwise height *p* < .001 (uncorrected) with clusters of at least 15 voxels retained. The dominant overlap cluster was located in the left lateral occipitotemporal extrastriate cortex, with a smaller overlap cluster in the left dorsal occipito-parietal/superior parietal region. Both clusters were located outside the independently defined AON search space except for one voxel.

**Table 1.**
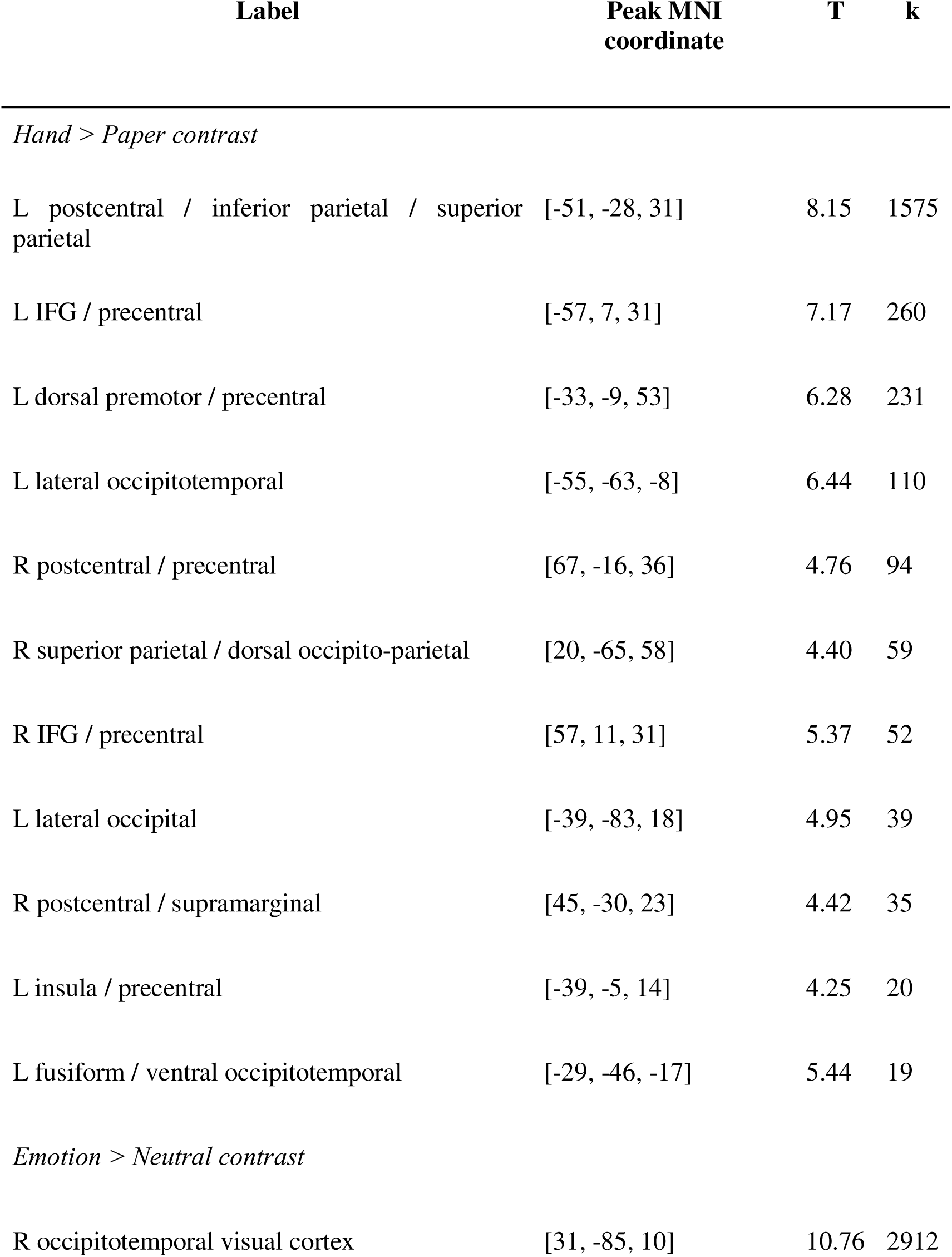
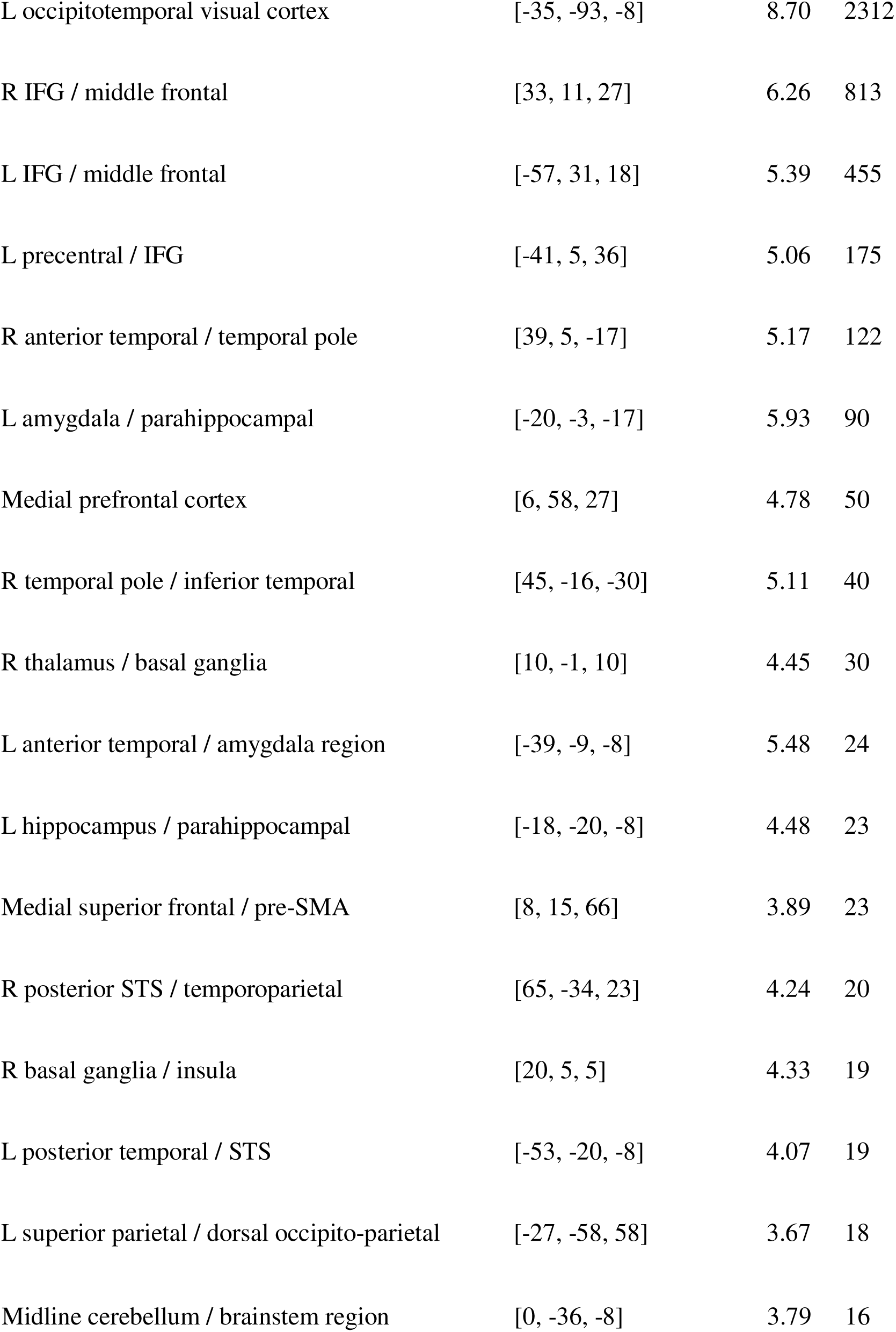
Whole-brain clusters for the hand action > paper control and emotional face > neutral face contrasts. Clusters were thresholded at voxelwise *p* < .001 uncorrected with k ≥ 15 voxels. Coordinates indicate peak MNI coordinates. T indicates peak t-value. k indicates cluster size in voxels.

| Label | Peak MNI coordinate | T | k |
| --- | --- | --- | --- |
| *Hand > Paper contrast* |  |  |  |
| L postcentral / inferior parietal / superior parietal | [-51, -28, 31] | 8.15 | 1575 |
| L IFG / precentral | [-57, 7, 31] | 7.17 | 260 |
| L dorsal premotor / precentral | [-33, -9, 53] | 6.28 | 231 |
| L lateral occipitotemporal | [-55, -63, -8] | 6.44 | 110 |
| R postcentral / precentral | [67, -16, 36] | 4.76 | 94 |
| R superior parietal / dorsal occipito-parietal | [20, -65, 58] | 4.40 | 59 |
| R IFG / precentral | [57, 11, 31] | 5.37 | 52 |
| L lateral occipital | [-39, -83, 18] | 4.95 | 39 |
| R postcentral / supramarginal | [45, -30, 23] | 4.42 | 35 |
| L insula / precentral | [-39, -5, 14] | 4.25 | 20 |
| L fusiform / ventral occipitotemporal | [-29, -46, -17] | 5.44 | 19 |
| *Emotion > Neutral contrast* |  |  |  |
| R occipitotemporal visual cortex | [31, -85, 10] | 10.76 | 2912 |
| L occipitotemporal visual cortex | [-35, -93, -8] | 8.70 | 2312 |
| R IFG / middle frontal | [33, 11, 27] | 6.26 | 813 |
| L IFG / middle frontal | [-57, 31, 18] | 5.39 | 455 |
| L precentral / IFG | [-41, 5, 36] | 5.06 | 175 |
| R anterior temporal / temporal pole | [39, 5, -17] | 5.17 | 122 |
| L amygdala / parahippocampal | [-20, -3, -17] | 5.93 | 90 |
| Medial prefrontal cortex | [6, 58, 27] | 4.78 | 50 |
| R temporal pole / inferior temporal | [45, -16, -30] | 5.11 | 40 |
| R thalamus / basal ganglia | [10, -1, 10] | 4.45 | 30 |
| L anterior temporal / amygdala region | [-39, -9, -8] | 5.48 | 24 |
| L hippocampus / parahippocampal | [-18, -20, -8] | 4.48 | 23 |
| Medial superior frontal / pre-SMA | [8, 15, 66] | 3.89 | 23 |
| R posterior STS / temporoparietal | [65, -34, 23] | 4.24 | 20 |
| R basal ganglia / insula | [20, 5, 5] | 4.33 | 19 |
| L posterior temporal / STS | [-53, -20, -8] | 4.07 | 19 |
| L superior parietal / dorsal occipito-parietal | [-27, -58, 58] | 3.67 | 18 |
| Midline cerebellum / brainstem region | [0, -36, -8] | 3.79 | 16 |

The thresholded intersection of the two maps contained 86 overlapping voxels. This overlap was almost entirely outside the independently defined AON search space, with 85 of the 86 overlapping voxels situated outside the AON union. The dominant overlap cluster (51 voxels, peak [-51, –58, –8]) was located in the left lateral occipitotemporal extrastriate cortex, near the lateral occipital/ventral occipitotemporal transition. A smaller additional overlap cluster was located in the left dorsal occipito-parietal/superior parietal region (16 voxels, peak [-24, –60, 58]). Thus, while whole-brain overlap was present, it was confined primarily outside the AON.

### Regional AON profiles show IFG co-engagement and posterior differentiation

A repeated-measures ANOVA on the ROI-average AON activation revealed a strong task × ROI-family interaction (*F*(2, 76) = 40.52, *p* < .001, η²_p_ = .516) (Figure 3). There was also a significant main effect of hemisphere (*F*(1, 38) = 14.33, *p* < .001, η²_p_ = .274) and significant interactions of task × hemisphere (*F*(1, 38) = 16.42, *p* < .001, η²_p_ = .302) and ROI-family × hemisphere (*F*(2, 76) = 11.60, *p* < .001, η²_p_ = .234). Neither the main effect of task (*F*(1, 38) = 3.12, *p* = .085, η²_p_ = .076), the main effect of ROI-family (*F*(2, 76) = 1.82, *p* = .170, η²_p_ = .046), nor the three-way task × ROI-family × hemisphere interaction was significant (*F*(2, 76) = 2.36, *p* = .101, η²_p_ = .058).

**Figure 3.**
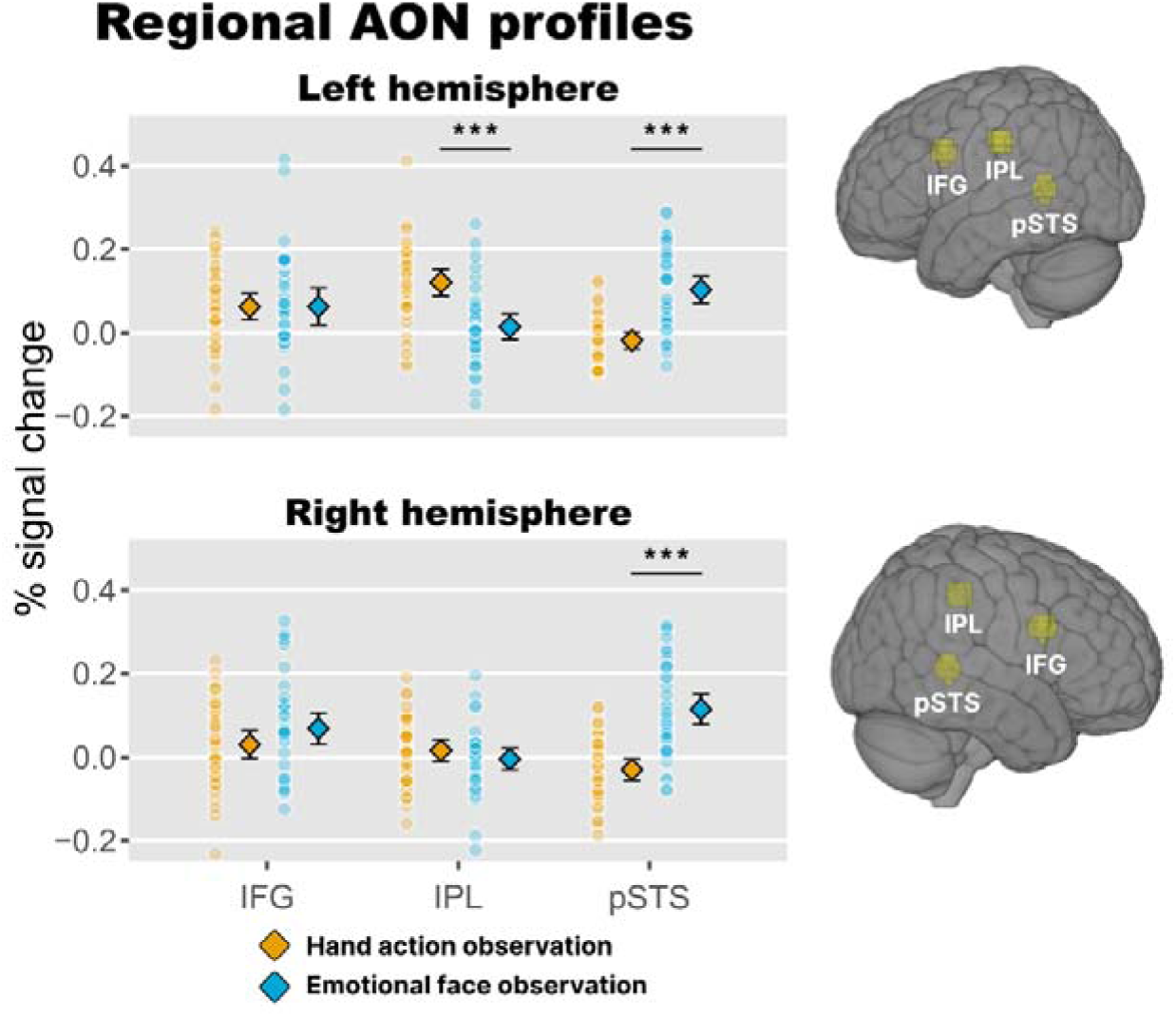
Regional AON response profiles. Percent signal change estimates are shown for bilateral IFG, IPL, and pSTS ROIs defined independently from meta-analytic action-observation coordinates (Caspers et al., 2010). Small points show individual participants, diamonds show group means, and error bars show 95% confidence intervals. The IFG showed positive responses to both contrasts without a reliable task difference, whereas the left IPL showed a hand action preference and bilateral pSTS showed an emotional face preference. ***: *p* < .001

Within the IFG, planned contrasts confirmed co-engagement, with positive responses for both hand action observation (mean = 0.0460, *t*(38) = 3.24, *p* = .00246, d = 0.52) and emotional face observation (mean = 0.0653, *t*(38) = 3.52, *p* = .00113, d = 0.56). The response did not differ between tasks at either the family level (mean difference = – 0.0193, *t*(38) = –0.77, FDR-corrected *p* = .447, d = –0.12) or the individual ROI level (left IFG: mean difference = –0.0003, *t*(38) = –0.01, corrected *p* = .991, d = 0.00; right IFG: mean difference = –0.038, *t*(38) = –1.34, corrected *p* = .283, d = –0.21).

In the posterior components, opposing task preferences emerged. The IPL family was more enhanced by hand action observation (mean difference = 0.0640, *t*(38) = 3.48, corrected *p* = .00191, d = 0.56), whereas the pSTS family was more activated in emotional face observation (mean difference = –0.1333, *t*(38) = –7.08, corrected *p* < .001, d = –1.13). The task × hemisphere interaction was explained by lateralisation differences: the IPL action preference was found only in the left hemisphere (left IPL: mean difference = 0.107, *t*(38) = 4.99, corrected *p* < .001, d = 0.80; right IPL: mean difference = 0.021, *t*(38) = 1.05, corrected *p* = .359, d = 0.17), whereas the pSTS emotion preference was present bilaterally (left pSTS: mean difference = –0.122, *t*(38) = –6.00, corrected *p* < .001, d = –0.96; right pSTS: mean difference = –0.145, *t*(38) = –6.48, corrected *p* < .001, d = –1.04).

### Voxelwise thresholded maps show minimal intersection within the AON

Within the AON, voxelwise thresholded maps for the two conditions were nearly non-overlapping (Figure 4A). At the height threshold of *p* < .001 (uncorrected), the AON union (709 voxels) contained 194 hand action voxels and 269 emotional face voxels, but only a single overlapping voxel (Dice coefficient = .0043). Hand action voxels occupied IFG (80 voxels) and IPL (114 voxels), whereas emotional face voxels were primarily found in pSTS (213 voxels) and in IFG (56 voxels), resulting in only one overlapping voxel (located in the left IFG, Dice coefficient = .0147) and zero in the posterior families. The absence of intersection shows that the activation maps remain almost segregated at the voxelwise scale, even within the IFG where regional co-engagement was observed at the ROI-average scale.

**Figure 4.**
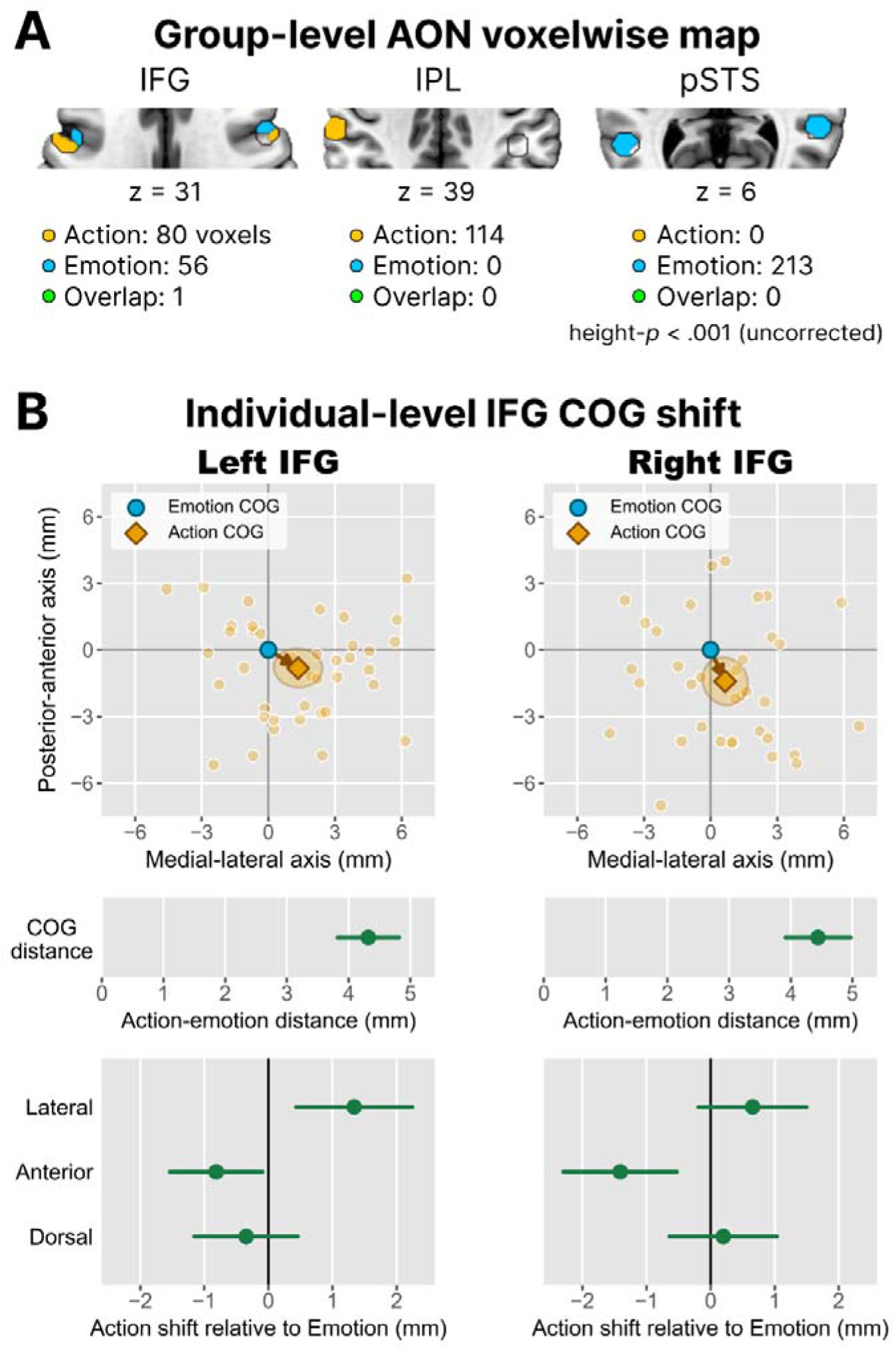
Voxelwise AON maps and individual-level IFG COG displacement. (A) Group-level voxelwise maps restricted to the independent AON search space. Maps were thresholded at voxelwise *p* < .001 uncorrected. Only one overlapping voxel was observed within the AON, located in the IFG, and no intersection overlap was observed in the posterior ROI families. (B) Individual-level t-weighted centre-of-gravity (COG) displacement within the IFG. Scatter plots show the individual hand action COG relative to the emotional face COG in the lateral-anterior plane. The orange diamond shows the group mean displacement. The shaded ellipse shows the 95% confidence region for the group mean displacement vector in the plotted two-dimensional plane. Lower plots show mean displacement components with bootstrap 95% confidence intervals. Positive values indicate hand action COG displacement in the lateral, anterior, or dorsal direction relative to the emotional face COG.

### Fine-grained spatial offset within the IFG

To investigate how regional co-engagement and voxelwise separation coexist, we evaluated individual-level spatial offsets within the bilateral IFG using t-weighted COGs (Figure 4B). In the left IFG, all 39 participants contributed paired COGs, yielding a mean Euclidean distance of 4.32 mm with 95% CI [3.86, 4.81], or 54% of the 8 mm ROI radius. The mean 3D offset vector differed reliably from zero (Hotelling *F*(3, 36) = 4.75, *p* = .0068). In the right IFG, 38 participants contributed paired COGs, yielding a mean distance of 4.44 [3.93, 4.96] mm, or 56% of the ROI radius (Figure 4B, lower). The mean 3D offset vector also differed reliably from zero (Hotelling *F*(3, 35) = 3.95, *p* = .0158).

The direction of these displacements along each MNI axis was partly consistent across hemispheres. In the left IFG, the hand action COG was situated significantly lateral to the emotional face COG by 1.34 [0.47, 2.20] mm (paired t-test, *t*(38) = 3.00, *p* = .0047, d = 0.48) and shifted posteriorly, as reflected by a negative displacement along the anterior axis, –0.82 [-1.51, –0.13] mm (*t*(38) = –2.29, *p* = .028, d = –0.37), but dorsal displacement was not (0.35 [-0.42, 1.11] mm, *t*(38) = 0.87, *p* = .387, d = 0.14). Similarly, in the right IFG, the hand action COG was significantly posterior by –1.41 [–2.26, –0.54] mm (*t*(37) = –3.23, *p* = .0026, d = –0.52). Lateral and dorsal displacement was not significant (0.65 [-0.16, 1.47] mm, *t*(37) = 1.57, *p* = .124, d = 0.26; –0.20 [-0.98, 0.62] mm, *t*(37) = –0.47, *p* = .638, d = –0.08). These results suggest that while both tasks recruit shared regions within the IFG, they exhibit a fine-grained, consistent posterior topographic bias, with clearer lateral separation in the left IFG, rather than a modular subdivision.

## Discussion

The present study demonstrates that the neural responses between hand action and emotional face observation are neither simply shared nor separate. Instead, their spatial overlap depends fundamentally on the scale of analysis. At the whole-brain level, overlap between the two contrasts was located to the left lateral occipitotemporal extrastriate cortex, outside AON boundaries. At the ROI-average scale, we confirmed regional co-engagement in the IFG, contrasted with the opposing task preferences of the IPL and pSTS. At the voxelwise scale, however, same-voxel intersection was virtually absent within the AON, with only one overlapping voxel located in the IFG. In addition, individual-level analysis revealed a reliable spatial offset of the IFG response centres between conditions. These findings suggest scale-dependent reuse, where the IFG is shared as a macro-anatomical region but not as a common same-voxel substrate.

The whole-brain maps clarify that the two contrasts exhibited some overlap outside the AON, as their dominant shared cluster was located in the left lateral occipitotemporal extrastriate cortex. This region contains adjacent or overlapping visual representations for body parts, faces, and biological motion (Haxby et al., 2000; Pitcher & Ungerleider, 2021). Recent fMRI evidence further supports its functional role that organises structured representations of human body parts into spatially proximate clusters for action effectors, such as hands, and facial parts (Kurihara et al., 2025). Such visual convergence is also consistent with the stimulus properties of our contrasts, given that the hand action task isolated a biological effector acting on objects, whereas the face task isolated expressive facial configurations. Of note, the lateral occipitotemporal cortex has also been shown to carry action information that generalises across observation and execution (Oosterhof et al., 2010). This overlap may therefore reflect convergence between social visual representations and action-related codes. A smaller overlap cluster was also observed in the left dorsal occipito-parietal and superior parietal region. This cluster may reflect a shared dorsal visual component for spatially guided, action-related visual processing (Culham & Valyear, 2006). Taken together, the overlap pattern points to convergence in occipitotemporal and dorsal visual–action regions outside the AON, rather than as evidence for a shared frontoparietal motor simulation mechanism.

Within the AON, the IFG exhibited regional co-engagement, showing positive responses to both hand action and emotional face observation at the ROI-average scale. This regional overlap is consistent with classic models of imitation and empathy that propose the IFG serves as a shared frontoparietal node for processing different social cues (Keysers & Gazzola, 2009; Rizzolatti & Craighero, 2004). However, macro-level regional overlap does not guarantee that different social domains recruit a common neural substrate. This is actually observed in recent social cognition studies, where apparent sharing in the human mirror system can be evaluated through shared voxel counts across social tasks and through directed interactions among STS, IPL, and IFG rather than through a single undifferentiated module (Sadeghi et al., 2022; Schmidt et al., 2021). Indeed, recent functional connectivity studies characterise the IFG as a heterogeneous region organised along continuous gradients (Diveica et al., 2023).

Consistent with this graded and individualised functional architecture, the current IFG co-engagement coincided with voxelwise displacement at individual-level. Between action and emotion observation, voxel intersection was virtually absent and a systematic offset was observed mainly along the posterior axis in the bilateral IFG, with a lateral shift in the left IFG. Rather than recruiting an identical, shared neural substrate, the two domains engage adjacent, potentially interdigitated representations within the same cortical region, mirroring a graded and locally heterogeneous IFG organisation (Diveica et al., 2023). Ferrari et al. (2017) proposed partially overlapping but distinct mirror pathways: a parietal–premotor sensorimotor pathway for hand actions and a face/mouth pathway with stronger limbic connections. This framework fits our finding that the two contrasts co-engaged the IFG but showed offset response centres. Relatedly, observed hand actions recruit premotor sectors dorsal to those recruited by observed mouth actions, forming a somatotopic pattern consistent with the motor homunculus (Buccino et al., 2001). Because the reliable shift in the present study was in the anteroposterior axis rather than dorsoventral, it may reflect a hand–face functional topography rather than a classical effector somatotopy. Thus, the IFG may be neither a uniform shared substrate nor two segregated modules, but rather a shared cortical region characterised by domain-biased local organisation. Although the present sensory observation design cannot test mirror-like motor properties directly, these spatial mapping results demonstrate that regional co-engagement masks a structured, fine-scale spatial segregation within the frontal AON.

The posterior AON nodes differentiated where the IFG co-engaged, following the within-task control contrasts. The left IPL responded to hand action relative to the paper control, but not to emotional faces relative to neutral faces. This fits evidence that anterior IPL represents object-directed action not in terms of physical trajectories or specific object forms, but abstract action meanings and goal structures (Caglar et al., 2025; Wurm & Erigüç, 2025). Conversely, the pSTS showed the opposite pattern, responding to emotional relative to neutral faces but not to hand action. The lack of a differential response in these nodes does not preclude their general involvement. In the IPL, subtracting neutral faces eliminates static form and identity, isolating emotional expressions. As these changeable affective cues are strongly associated with pSTS-centred expression processing (Pitcher & Ungerleider, 2021), the IPL reasonably remains unresponsive. For the pSTS, the lack of response to hand actions does not imply insensitivity to observed actions (Karakose-Akbiyik et al., 2023). Rather, the subtraction of the paper control removes shared spatial kinematics, such as the approach trajectory and contact event, which may cancel out the motion-driven pSTS response. Thus, posterior nodes tracked the specific sensory-motor properties of their preferred inputs, whereas the IFG engaged across both formats despite minimal voxelwise overlap.

A main limitation lies in the fixed task order. The motor observation task always preceded the emotional face task to prevent emotional stimuli from pre-activating frontal simulation systems (Del Vecchio et al., 2024). This fixed order, however, confounds task domain with time and fatigue effects, restricting our conclusions to spatial mapping within specific contrasts rather than a general action-versus-emotion dissociation. Furthermore, the stimulus formats differed between dynamic hand videos and static face images. The two contrasts were also not strictly matched for detection demand, as target detection was slower for hand actions than for paper controls and for emotional than for neutral faces. Such differences in attentional demand could have produced the regional responses we observed, although they are unlikely to account for the systematic spatial offset between the two response centres within the IFG. Future studies should match stimulus formats and detection demand, and include execution or mimicry conditions to directly test the mirror-like coding of emotional expressions (Sato et al., 2023; Wallenwein et al., 2024).

In summary, whole-brain convergence was concentrated in the left lateral occipitotemporal cortex, while the IFG showed co-engagement at the ROI-average scale but minimal voxelwise overlap and a systematic posterior bias. The IPL and pSTS retained opposing contrast preferences. Together, these findings show how regional reuse can coexist with local specialisation: the same cortical region can support responses to hand actions and emotional faces without erasing the topographic structure that distinguishes them.

## Author Notes

### Data availability

Data and analysis code are available at https://osf.io/r58jx. Video clips used in the motor observation task are also available on request.

### Constraints on generality

The present sample consisted of right-handed young Japanese adults. The conclusions are intended to generalise to the spatial relationship between these two task contrasts under passive observation with attention checks. They should not be assumed to generalise to other age groups, left-handed participants, clinical populations, dynamic emotional expressions, action execution, imitation, or designs in which task order and stimulus format are fully counterbalanced.

